# Air Pollution Aggravates Renal Ischemia-Reperfusion-Induced Acute Kidney Injury

**DOI:** 10.1101/2023.08.17.553734

**Authors:** Talita Rojas Sanches, Antonio Carlos Parra, Peiqi Sun, Mariana Graner, Lucas Yuji Umesaki Itto, Loes M. Butter, Nike Claessen, Joris J.T.H. Roelofs, Sandrine Florquin, Mariana Matera Veras, Maria de Fátima Andrade, Paulo Hilário Nascimento Saldiva, Jesper Kers, Lucia Andrade, Alessandra Tammaro

## Abstract

Chronic kidney disease (CKD) has emerged as a significant global public health concern. Recent epidemiological studies have highlighted the link between exposure to fine particulate matter (PM2.5) and declined renal function. PM2.5 exerts its harmful effects on various organs through oxidative stress and inflammation. Acute kidney injury (AKI) resulting from ischemia reperfusion injury (IRI) involves similar biological processes involved in PM2.5 toxicity and is a known risk factor for CKD. The objective of this study was to investigate the impact of PM2.5 exposure on IRI-induced AKI. Mice were exposed to PM2.5 or filtered air for 12 weeks before IRI, and were euthanized 48h after IRI. Animals exposed to PM2.5 and IRI exhibited reduced glomerular filtration and impaired urine concentration ability. Moreover, they showed elevated tubular damage markers NGAL and KIM-1, along with significant tubular necrosis. PM2.5 exposure exacerbated local innate immune activation, leading to an increased infiltration of Ly6G+ granulocytes and F480+ macrophages in the kidney. This, in turn, contributed to heightened renal senescence markers and myofibroblast infiltration. Collectively, our findings suggest that AKI-induced hampered tubular function is worsened by PM2.5, leading to reduced resilience to stress, activation of aging mechanisms and early hallmarks of fibrosis. Decreasing PM2.5 and implementing preventive strategies can improve AKI patients outcome and prevent AKI progression.

## INTRODUCTION

Air pollution is a well-recognized environmental health risk^1^. Particulate matter (PM), brings public health concerns because of its toxicity and the ubiquitous human exposure to this pollutant^2^. PM, which includes inhalable particles with an aerodynamic diameter of 10 μm or less (PM10) and fine particles with an aerodynamic diameter of 2.5 μm or less (PM2.5), is emitted from combustion sources (light vehicles and heavy-duty diesel vehicles) or formed through atmospheric chemical transformation (gas-to-particle conversion)^3^. Environmental levels of PM2.5 serve as air quality indicator.

A significant body of evidence links short and long-term PM2.5 exposure to negative effects on the respiratory system^4,5,6^, cardiovascular system^7,8^, brain function^9,10^ and cancer^11^. Recently, Liu and colleagues demonstrated dependent associations between short-term exposure to PM10 and PM2.5 and daily all-cause, cardiovascular, and respiratory mortality in more than 600 cities across the globe^12,13^. Until 2018 the effect of PM2.5 on kidney function was largely neglected^14^. Since then, epidemiological evidences demonstrated that long-term exposure to PM2.5 is associated with an increased risk of incident Chronic Kidney disease (CKD)^15,16,17^, CKD progression^16^, albuminuria^18^, end stage kidney disease (ESKD)^14^ and post-transplant outcomes^19^. Epidemiological studies have the limitation of being unable to assess accurately individual exposure to PM2.5.

Zip-code levels of PM2.5 are commonly used to determine individual exposure but actual exposure to PM2.5 could be influenced by occupational activity, mobility patterns, residence’s proximity to air pollution sources and indoor air pollution^20^. Therefore, environmentally-controlled experimental studies are needed to bypass these confounders and to elucidate the renal pathological mechanisms initiated by PM2.5.

Few studies have described kidney damage induced by PM2.5 exposure in experimental models with contrasting data about its effect on inflammation and toxicity^21,22^. Oxidative stress induction is a central paradigm for the pro-inflammatory effect of PM2.5 exposure^23^, but also a central mechanism of damage during an episode of acute kidney injury (AKI)^24^. AKI can occur during renal transplant procedures, hypotension, all causes of shock, or due to sepsis and is one of the most common risk factors for CKD progression^25^. In this study, we aim to elucidate the impact of PM2.5 in an experimental model of renal ischemia-reperfusion (IRI)-induced AKI.

For PM2.5-controlled exposure we used a unique equipment, the Harvard Ambient Particle Concentrator^26^, where the exposure is performed in whole-body chambers. Mice received PM2.5-rich or filtered air for 12 weeks before the induction of IRI. Next, we analyzed kidney function and performed an in-depth molecular and histological analysis of the kidney. The results of this study shed light on mechanisms worsened by PM2.5 after induction of AKI. Moreover, it highlights previously unknown pathological pathways that can be targeted for prevention, ultimately reducing the burden of kidney disease.

## METHODS

### Mice

All experimental procedures carried out in this study have been approved by the Medical Research Ethics Committee of the University of São Paulo School of Medicine (Animal Ethics Committee, No. 1205/2019) and were conducted in accordance with the National Institutes of Health Guide for the Care and Use of Laboratory Animals. Male six-week-old C57Bl/6 animals (20-25 g body weight) were obtained from the animal facility of the University of São Paulo School of Medicine. Mice were kept under standard environmental conditions (temperature, humidity, ventilation, light/dark cycle), housed in specific pathogen-free conditions (SPF) with *ad libitum* access to water and food.

### Air pollution exposure model

Animal exposure to PM2.5 was conducted using the Harvard Ambient Particle Concentrator (HAPC), a widely utilized method for controlled exposure of rodents to air pollution^27^. Figure 1A illustrates the functioning of HAPC, which concentrates ambient levels of PM2.5 by employing a series of virtual impactors that selectively allow the passage of particles smaller than 2.5 micrometers. The concentrated stream of PM2.5 is then directed into the exposure chamber at a flow rate of 10 liters per minute. HAPC concentrates ambient PM2.5 levels by a factor 20. For instance, if the ambient PM2.5 level is 10 μg/m^3^, HAPC increases it to 200 μg/m^3^. Consequently, to achieve a cumulative dose of 600 μg/m3 (equivalent to the daily exposure levels experienced by individuals living in highly polluted cities like São Paulo and many others), the duration of exposure varied based on the daily ambient PM2.5 levels. The formula used to determine the exposure time in the HAPC was as follows:

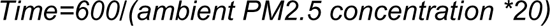

**Figure 1.**
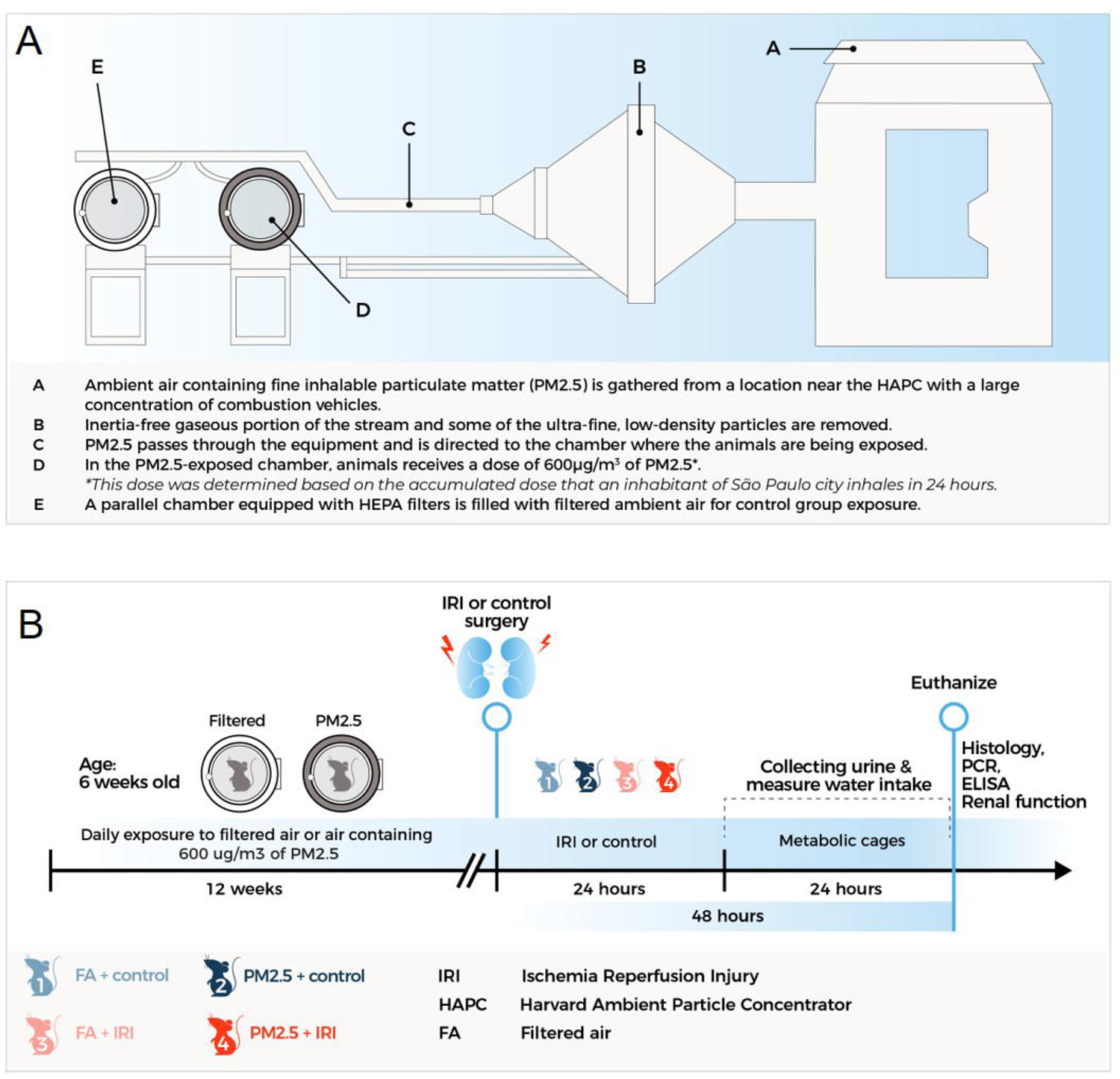
HAPC function and experimental setup. (A) Air with PM 2.5 is gathered from around HPAC location, a place with large concentration of combustion vehicles (A); Inertia-free gaseous portion of the stream and some of the ultra-fine, low density particles are removed (B); Fine inhalable particulate matter (PM 2.5) passes through the equipment and is directed to the animals being exposed (C); In the exposed chamber, animals were exposed daily to 600 ug/m3 of PM 2.5. This is the equivalent of what an individual living in the metropolitan area of São Paulo receives daily (D); A parallel chamber is fed with filtered ambient air for the control group exposure(E). (B) Experimental setup: Six weeks-old mice were exposure to filtered air or polluted air (PM 2.5) for 12 weeks. Some of them were subjected to bilateral renal ischemia for 30 min followed by reperfusion. Control and PM 2.5 mice were evaluated 48h after IRI.

In the control group, animals were exposed to filtered air (FA) under identical conditions. A separate chamber equipped with high-efficiency particulate air (HEPA) filters was used to effectively remove PM2.5 from the air. Strict control over temperature and humidity conditions was maintained within the chambers, following the protocols established by the Experimental Atmospheric Pollution Laboratory^28,29^. These experiments complied with standard protocols. The HAPC is located in the garden of the University of São Paulo School of Medicine (23°33’19”S; 46°40’22”W), near a major intersection with heavy vehicular traffic. Consequently, the mice were exposed to PM2.5 originating from vehicle emissions.

### Surgical model of renal IRI

After 12 weeks of exposure to PM2.5 or FA, mice were subjected to the surgical model of renal IRI. Briefly, mice were anesthetized with ketamine (90 mg/kg body weight) and xylazine (10 mg/kg body weight). Subsequently, through a midline abdominal incision, kidneys were exposed and both renal arteries were clamped for 30 min (ischemia). Afterwards, clamps were removed to restore blood circulation (reperfusion). The entire procedure was performed on a heated pad for temperature-controlled surgery. After IRI, mice were allowed to recover for about one hour in isolated cages, kept on the heated pad to maintain constant body temperatures. When animals became active again, we gave a subcutaneous injection of morphine for analgesic purposes (10mg/kg BW). During the 24-hours recovery period, drinking water was supplemented with dipyrone (200 mg/kg BW). Twenty-four hours after surgery, mice were moved to metabolic cages, where they remained for an additional 24 h, for the collection of urine over a 24-hour period. During this period, water intake was recorded, and urine volume was measured. Animals were euthanized 48 h post-IRI by receiving an intraperitoneal injection of high doses of ketamine (90 mg/kg body weight) and xylazine (10 mg/kg body weight), followed by heart puncture for blood collection. Kidneys were harvested for different analysis. The sample size of the different groups was as follow: FA+control (n=4), PM2.5+control (n=7), FA+IRI (n=11), PM2.5+IRI (n=15).

### Biochemical analysis

Urine and blood samples were aliquoted and centrifuged for 30 minutes at 4.000g. Quantification of serum and urine sodium levels was performed using an autoanalyzer (EasyLyte; Medica Corporation). Creatinine concentration was determined using a commercial kit (Creatinine kit; Labtest Diagnostic). Creatinine clearance was calculated with the following formula:

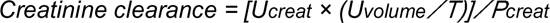

where Ucreat is the urinary concentration of creatinine (in mg/dL), Uvolume is the urine volume (in µL), T is the time (in min), and Pcreat is the plasma concentration of creatinine (in mg/dL). We measured urine osmolality using a freezing-point osmometer (3D3; Advanced Instruments).

### (Immuno)#histochemistry

Kidneys were fixed in formalin for 24hrs, embedded in paraffin and cut in 4-µm thick sections for histology. For tubular necrosis score, paraffin sections were stained using Periodic acid-Schiff diastase method to assess the extent of tubular injury, characterized by loss of renal brush border, presence of protein casts, as well as presence of necrosis and tubular dilation. Quantification of tubular necrosis was assessed by 2 pathologists (S.F. and J.J.R) in a blinded fashion. Briefly, we adopted a five-point scale to classify the proportion of corticomedullary region involved in 10 non-overlapping high-power fields (×100 magnification), according to the following criteria^30^: 0 = absent; 1 = 0-10%; 2 = 10-25%; 3 = 25-50%; 4 = 50-75%; and 5 = 75-100%.

For immunohistochemistry: slides were deparaffinized, rehydrated and incubated in 0,3% H_2_O_2_ in methanol to block endogenous peroxidase. Antigen retrieval was performed by pepsin digestion for 30 minutes (0,25% in 0,01M HCl) for Ly6G or by boiling in 0,01M citrate buffer (pH 6,0) for 10 minutes at 120°C for all other antigens. Antibodies were diluted in antibody diluent (Immunologic) and slides were incubated overnight at 4°C with the following antibodies: rat anti-mouse F4/80 (Serotec, MCA497), rabbit anti-mouse p21 (Abcam, ab188224), rat anti-mouse Klotho (Transgenic, KO603), mouse IgG2a anti-mouse αSMA (Dako, M851), rabbit anti-mouse vimentin (Cell Signaling, 5471) and anti-mouse Ly6G-FITC (BD, 553126). As secondary antibodies, goat anti-mouse IgG2a HRP (Southern Biotech), rabbit anti-rat (Southern Biotech), rabbit anti-FITC (Dako) and Brightvision goat anti-rabbit HRP (Immunologic) were used. Incubation was done for 30 minutes at room temperature. BrightDAB (Immunologic) was used as a substrate to detect HRP.

### ELISA and western blot

Urinary neutrophil gelatinase-associated lipocalin (NGAL) was measured by using a commercially available ELISA kit (R&D Systems), according to the instructions provided by the manufacturer. For western blot, kidney lysates were prepared from 10 frozen sections (20 µm thick) incubated at 4°C for 30 min in RIPA buffer containing 50 mM Tris pH7.5, 0.15 M NaCl, 2 mM EDTA, 1% deoxycholic acid, 1% NP-40, 4 mM sodium orthovanadate, 10 mM sodium fluoride, 1% protease inhibitor cocktail (P8340, Sigma). The lysates were then centrifuged at 12000 rpm for 15 minutes and the supernatants were collected and stored at −20°C. SDS-polyacrylamide gel electrophoresis was carried out on precasted 4-12% gels (Thermo Fisher Scientific) and proteins were electrophoretically transferred onto methanol-activated polyvinylidene fluoride (PVDF) microporous membranes (Millipore). Membranes were blocked for one hour with 5% bovine serum albumin (Sigma) in Tris-buffered saline containing 0.1% Tween 20 (TBS-T), followed by overnight incubation at 4uC with primary rabbit anti-mouse/human STING (Cell Signaling #13647), cGAS (Cell signaling #31659), pIRF3 (Cell signaling #29047), IRF3 (Abcam #68481) and β-actin (Sigma Merck A5441). HRP-conjugated secondary antibodies (DAKO) were incubated for one hour at room temperature, and HRP activity was visualized with ECL-reagent. β-actin was used as loading control. Densitometric quantification analysis was performed on imagines of scanned films using ImageJ.

### Quantitative Real Time-PCR

Total RNA was extracted from the frozen kidney section with Trizol reagent (Invitrogen) according to the manufacturer’s protocol. cDNA was synthesized using M-MLV transcriptase (Promega) and oligo dT primers. Quantitative RT-PCR was performed on a LightCycler480 (Roche) using SensiFAST SYBR NO ROX kit (Bioline). Fluorescent dye intensity was analyzed and quantified with linear regression. Gene expression was normalized against the housekeeping gene, TATA box binding protein (TBP). The sequence of primers used is as follows: *Tbp* se: ggagaatcatggaccagaaca as: gatgggaattccaggagtca; *Cxcl1* se:ataatgggcttttacattctttaacc as: agtcctttgaacgtctctgtcc; *Ccl2* se: catccacgtgttggctca as: gatcatcttgctggtgaatgagt; *Il1b* se: tgagcaccttcttttccttca as: ttgtctaatgggaacgtcacac; *Cxcl2* se: ccctggttcagaaaatcatcc as: cttccgttgagggacagc; *Tgfb* se: gcaacatgtggaactctagaa as: gacgtcaaaagacagccactca; *Pai1* se: aagtctttccgaccaagag as: ctgagatgacaaaggctgtg; *Kim1* se: tggttgccttccgtgtctct as: tcagctcgggaatgcacaa; *Cxcl10* se: gctgccgtcattttctgc as: tctcactggcccgtcatc; *Ifnb1* se: ggaaagattgacgtgggaga as:cctttgcaccctccagtaat; *Pdgf* se: agtgatgtctggtcttttggg as: tggcattgtagaactggtcg

### Concentrations and trace-elements composition of PM2.5

Urban atmospheric PM2.5 aerosol and trace elements were analyzed over the months during which the animals were exposed (from 07/11/19 to 12/18/19). The membrane filters collected were sampled and analyzed for the identification of trace elements and determination of their masses by inductively coupled plasma mass spectrometry (ICPMS) or ion chromatography. The PM2.5 mass concentrations were determined by gravimetric analysis of the Teflon filters.

### Statistical analysis

All data are presented as mean ± standard error of the mean (SEM). Statistical analyses were performed using a one-way analysis of variance (ANOVA), followed by the Kruskal-Wallis test. The non-parametric Mann Whitney test was used for 2 groups comparison. Statistical significance was determined by a P-value of or less than 0.05. The statistical analysis was performed using GraphPad Prism software, version 8.0 (GraphPad Software).

## RESULTS

### Experimental design and full description of particle concentration and elemental composition

To facilitate the understanding of our experimental design we added in Figure 1A-B a visual representation of the exposure system (Fig. 1A), timeline and the experimental groups (Fig. 1B). Mice were exposed in the HAPC from July until September 2019. Supplemental Table 1 provides a comprehensive and detailed depiction of both the chemical composition and concentration of PM2.5 throughout the entire exposure period. The ambient air concentration of PM2.5 during the experiment was 29.8±11 ug/m^3^. The elemental composition analysis revealed a high concentration of Organic Carbon (OC) and Elemental Carbon (EC), followed by metals. These elements, particularly elemental carbon and metals are directly emitted from combustion processes. These findings align with earlier studies that indicated vehicular emission as the main source of PM2.5 in the Metropolitan area of São Paulo ^3^.

### PM2.5 aggravates tubular damage in IRI-induced AKI

In order to investigate the impact of PM2.5 on IRI-induced AKI, we first looked at glomerular and tubular function. A sharp decline in creatinine clearance (used as a proxy for the glomerular filtration rate) was detected in the PM2.5+IRI group compared with FA+IRI (Fig. 2A) whilst no differences were detected among other groups. Furthermore, we observed a notable decrease in urinary osmolality as shown in Fig. 2A, indicating that PM2.5 plays a crucial role in impairing the kidney’s ability to concentrate after IRI. This was also supported by an increased fraction excretion of sodium, also a marker of tubular dysfunction (Fig. 2A). Therefore, we measured biomarkers of tubular damage. Urinary NGAL and renal KIM-1 gene expression were higher in the PM2.5+IRI group than in the FA+IRI group (Fig. 2B). Finally, renal pathology revealed a sharp increase in tubular damage in the PM2.5+IR group compared to FA+IRI, detected by PAS-D and expressed as tubular necrosis score (Fig. 2C).

**Figure 2.**
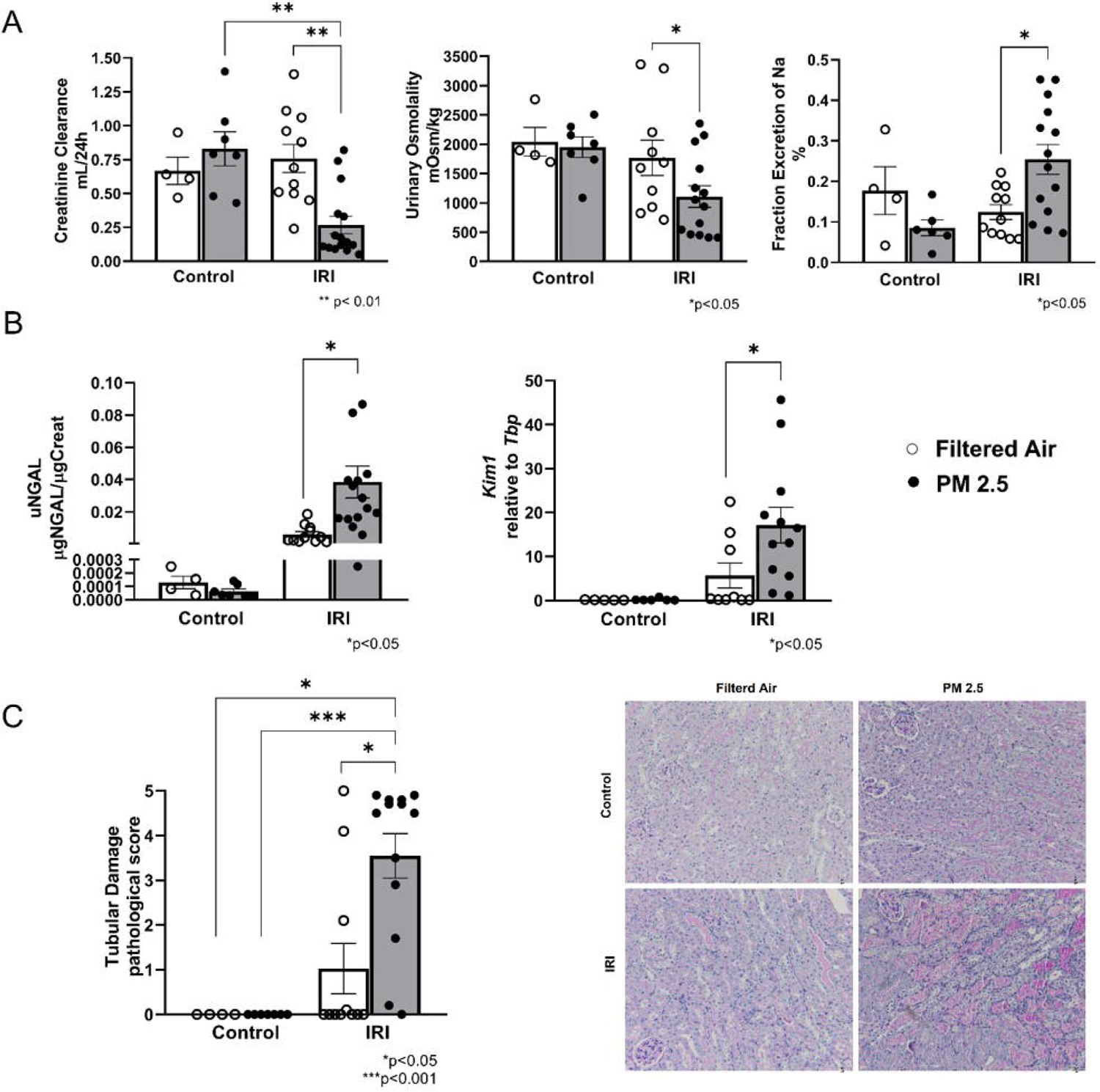
PM2.5 exacerbates IRI-induced kidney dysfunction. (A) Renal function was assessed by determining creatinine clearance with the formula: creatinine clearance = (U_creatinine_ × U_volume_/1,440 min)/P_creatinine_, where U_creatinine_ is urinary concentration of creatinine, U_volume_/1,440 min is urine output in microliters per day (1,440 min), and P_creatinine_ is plasmatic concentration of creatinine. (Statistical Analysis: Kruskal-Wallis test multi comparisons between all groups); Urinary Osmolality concentration was assessed by freezing-point osmometer (Statistical Analysis: Kruskal-Wallis test multi comparison between FA+IRI and PM2.5+IRI); Urinary and plasmatic sodium concentration was assessed by selective ion electrode method. The fraction of Na+ was calculated with the formula: (U_Na_ x U_creatinine_) / (P_Na_ x P_creatinine_) x 100, where UNa is urinary concentration of sodium, U_creatinine_ is urinary concentration of creatinine, PNa is plasmatic concentration of sodium and P_creatinine_ is plasmatic concentration of creatinine (Statistical Analysis: Kruskal-Wallis test multi comparison between FA-IRI and PM2.5-IRI). (B) Concentration of urinary NGAL measured by ELISA (Statistical Analysis: Kruskal-Wallis test multi comparison between FA+IRI and PM2.5+IRI); gene expression of *Kim1* in renal tissue assessed by real-time PCR (Statistical Analysis: Kruskal-Wallis test multi comparison between FA+IRI and PM2.5+IRI) (C) Tubular injury score assessed on kidney tissue sections stained with periodic acid-Schiff (Statistical Analysis: Kruskal-Wallis test multi comparisons between all groups). High resolution images of PAS staining. White dot/ White bar: Filtered Air; Black dot/Gray bar: PM2.5.

### PM2.5 enhances renal inflammation via cGAS-STING

Renal tubular damage drives a local inflammatory response attracting the infiltration of innate immune cells into the renal parenchyma^31^. Renal chemokines and cytokines production function as chemoattractants for immune cells. Gene expression levels of renal pro-inflammatory chemokines Cxcl1*, Ccl2, Cxcl2* and cytokine *Il1b* were increased in animals subjected to IRI breathing PM2.5 (Fig. 3A). Consequently, the number of Ly6G-positive inflammatory granulocytes infiltrating the renal tissue was significantly higher in the PM2.5+IRI group compared with FA+IRI, but also in the IRI groups compared to control groups (Fig. 3B). F4/80+ macrophage infiltration exhibited a similar pattern, indicating that PM2.5 exacerbates renal inflammation (Fig. 3B). We then investigated whether the enhanced inflammatory response caused by PM2.5 could be linked to the activation of the cyclic GMP-AMP synthase (cGAS) stimulator of interferon genes (STING) pathway. This pathway is triggered by tubular mitochondria damage during AKI and leads to inflammation and AKI progression^32^. Additionally, in cultured TECs and in many other cells, PM2.5 was also shown to induce mitochondrial damage^33–35^. Strikingly, we found that the PM2.5+IRI group showed a significant activation of the cGAS-STING signaling pathway, including the phosphorylation of Interferon regulatory factor 3 (pIRF3) (Fig. 3C), which promotes the transcription of IFNß and NF-LB. The transcription of both factors was corroborated by the enhanced expression of the *Ifnb* and *Cxcl10* genes, downstream of both pathways respectively (Fig. 3C). Altogether this suggests that PM2.5 and IRI act synergistically to directly enhance inflammation by activating the cGAS-STING pathway, possibly via a compromised mitochondrial integrity in the injured tubular epithelium.

**Figure 3.**
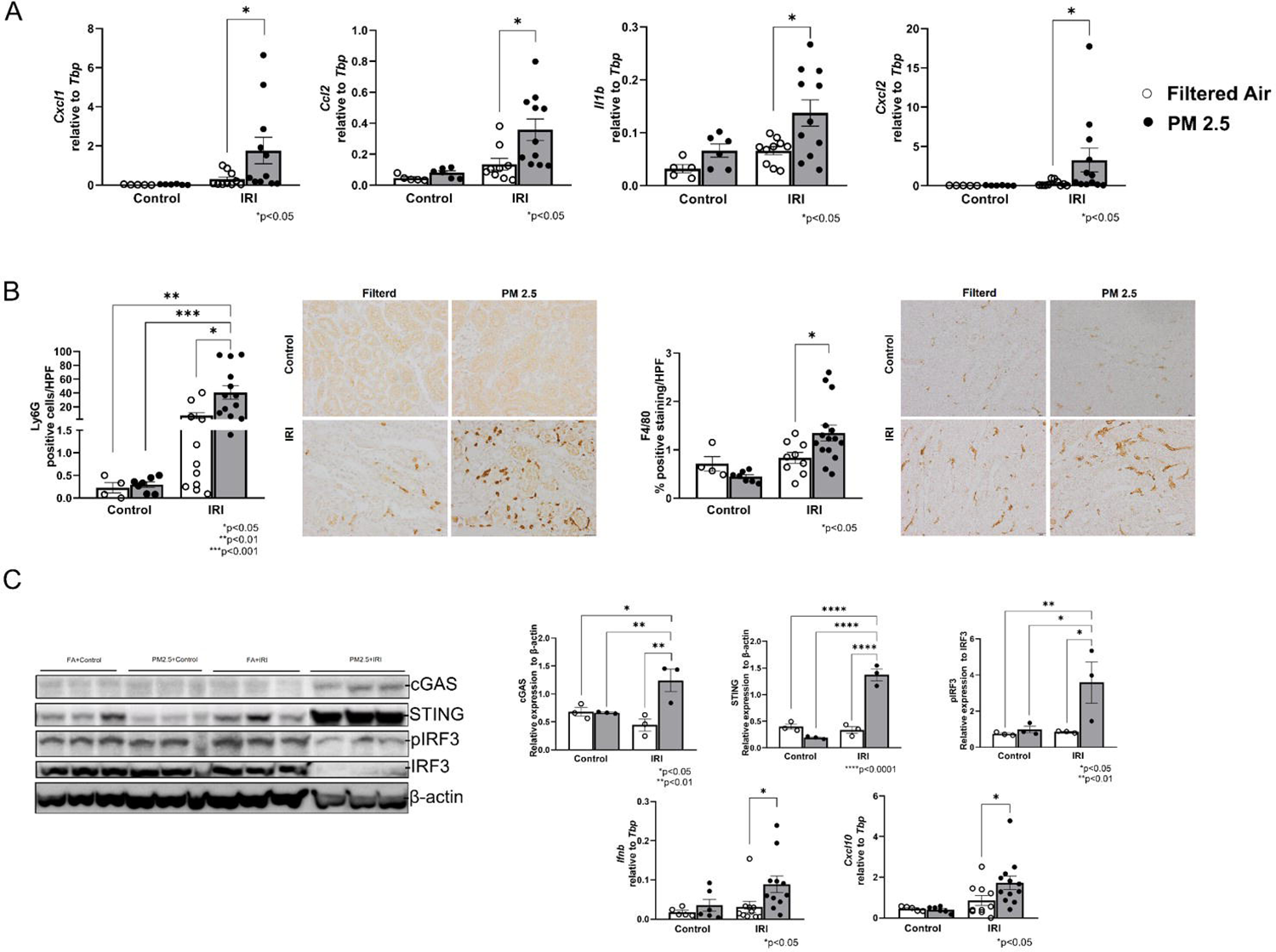
PM2.5 enhances renal inflammation. (A) Gene expression of *Cxcl1, Ccl2, Il1b* and *Cxcl2* in renal tissue measured by RT-PCR relative to *Tbp* (Statistical Analysis: Mann-Whitney test between FA-IRI and PM2.5-IRI, for *Cxcl1*; Kruskal-Wallis test multi comparison between FA-IRI and PM2.5-IRI, for *Ccl2*; Mann-Whiteny test between FA-IRI and PM2.5-IRI, for *Il1b*; Mann-Whitney test between FA-IRI and PM2.5-IRI, for *Cxcl2*). (B) Quantification and representative images of kidney tissue sections stained with Ly6G antibody showing granulocyte infiltration (Statistical Analysis: Kruskal-Wallis test multi comparisons between all groups). Quantification and representative images of kidney tissue sections stained with F4/80 antibody showing macrophage infiltration (Statistical Analysis: Mann-Whitney test between FA-IRI and PM2.5-IRI). (C) Immunoblotting of cGAS, STING, pIRF3, IRF3 and β-actin protein expression in renal tissue (Statistical Analysis: One Way ANOVA test multi comparisons between all groups); Gene expression of *Ifnb* and *Cxcl10* in renal tissue measured by RT-PCR (Statistical Analysis: Kruskal-Wallis test multi comparisons between all groups, for *Ifnb*; Kruskal-Wallis test multi comparison between FA-IRI and PM2.5-IRI, for *Cxcl10*). White dot/ White bar– Filtered Air; Black dot/Gray bar – PM2.5.

### PM2.5 accelerates aging mechanisms and triggers early hallmarks of fibrosis

Previous studies have demonstrated that PM2.5 can accelerate aging via different mechanisms^36^, among which oxidative stress^37,38^, inflammation^2^, DNA damage^39^ and cellular senescence^33,40^. Considering the high extent of renal damage and that cGAS-STING-mediated inflammation is linked to progression after AKI, we questioned whether IRI and PM2.5 could act synergistically to accelerate renal aging. Accumulation of p21+ senescent cells is one of the main mechanisms driving maladaptive repair after AKI leading to premature aging^41^. Renal *Cdkn1a* gene expression and p21+ cells detected by IHC, were significantly increased in the PM2.5+IRI group compared with PM2.5+Control and FA+IRI groups (Fig. 4A). This was also associated with an increased expression of genes associated with the senescence-associated secretory phenotype (SASP), a mechanism by which senescence cells exert their detrimental effect when abundant in renal tissue^42^. Renal gene expression of pro-fibrotic SASP components *Tgfb, Pdgf* and *Pai1* is increased in PM2.5+IRI compared to FA+IRI (Fig.4B). Additionally, during premature aging, there is a loss of renal protective mechanisms, such as the anti-aging factor Klotho^43^. We found that IRI induced a decline in klotho protein expression compared to control mice, but this decline was even more profound in the group breathing PM2.5 (Fig. 4C).

**Figure 4.**
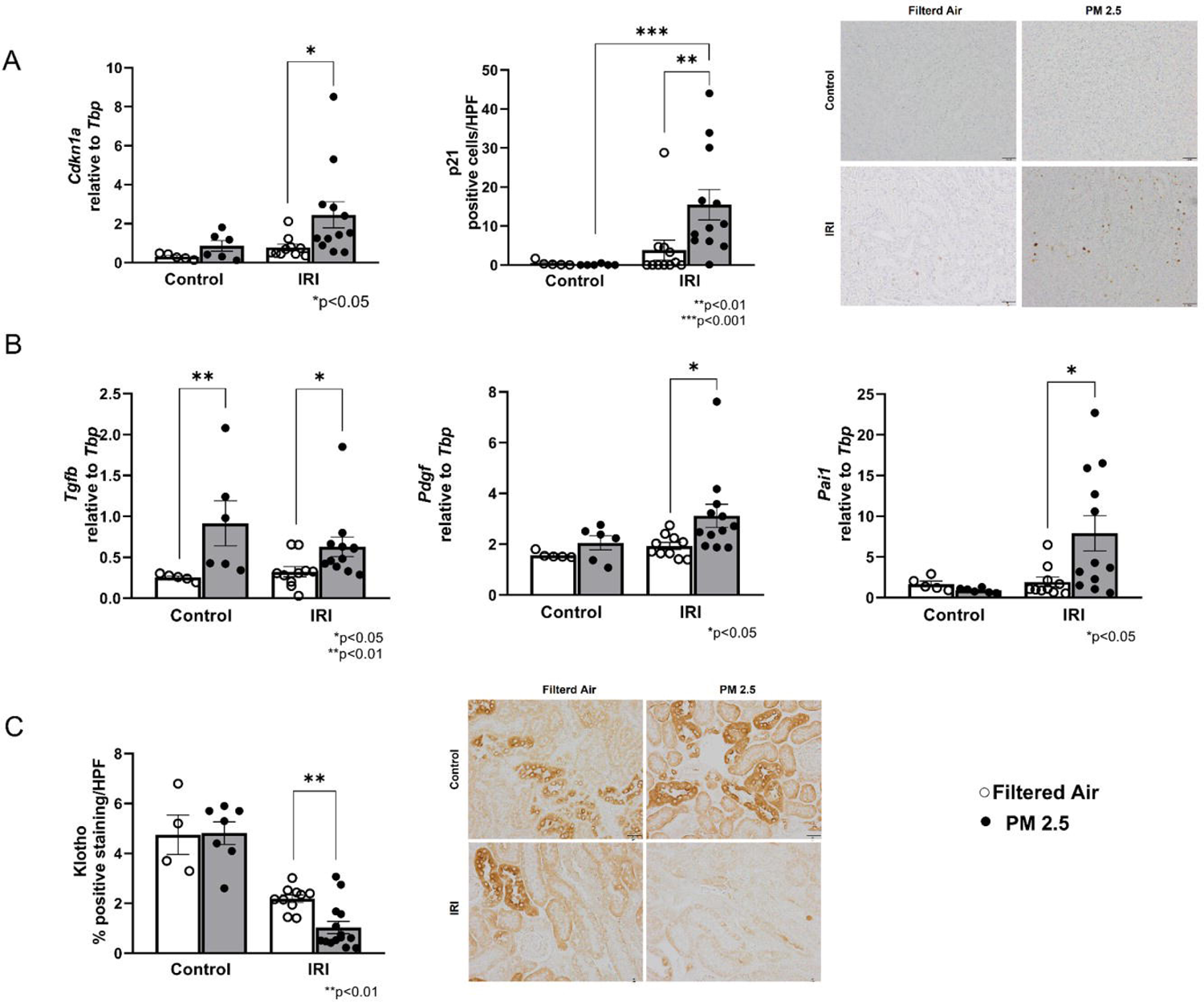
Accelerated aging mechanisms driven by PM2.5. (A) Gene expression of *Cdkn 1a* in renal tissue measured by RT-PCR relative to *Tbp*. Quantification and representative images of kidney sections stained with p21 antibody (Statistical Analysis: Kruskal-Wallis test multi comparisons between all groups); (B) Gene expression of *Tgfb, Pdgf* and *Pai1* in renal tissue measured by RT-PCR relative to *Tbp* (Statistical Analysis: One Way Anova follow by Kruskal-Wallis test multi comparison between FA-IRI and PM2.5-IRI and between FA-Sham and PM2.5-Sham, for *Tgfb*; Mann-Whiteny test between FA-IRI and PM2.5-IRI, for *Pai1;* Kruskal-Wallis test multi comparison between FA-IRI and PM2.5-IRI, for *Pdgf*); (C) Quantification and representative images of kidney sections stained with Klotho antibody (Statistical Analysis: Mann-Whiteny test between FA-IRI and PM2.5-IRI). White dot/ White bar– Filtered Air; Black dot/Gray bar – PM2.5

We then sought to assess whether these changes could trigger early hallmarks of fibrosis, associated with AKI progression^24^. Mice exposed to PM2.5+IRI show increased presence of α-smooth muscle actin (SMA) positive interstitial myofibroblasts, and vimentin expression compared with the FA counterparts (Fig 5A-B). Taken together our results demonstrated that PM2.5 triggers molecular mechanisms associated with premature aging and initiates early processes preceding renal fibrosis.

**Figure 5.**
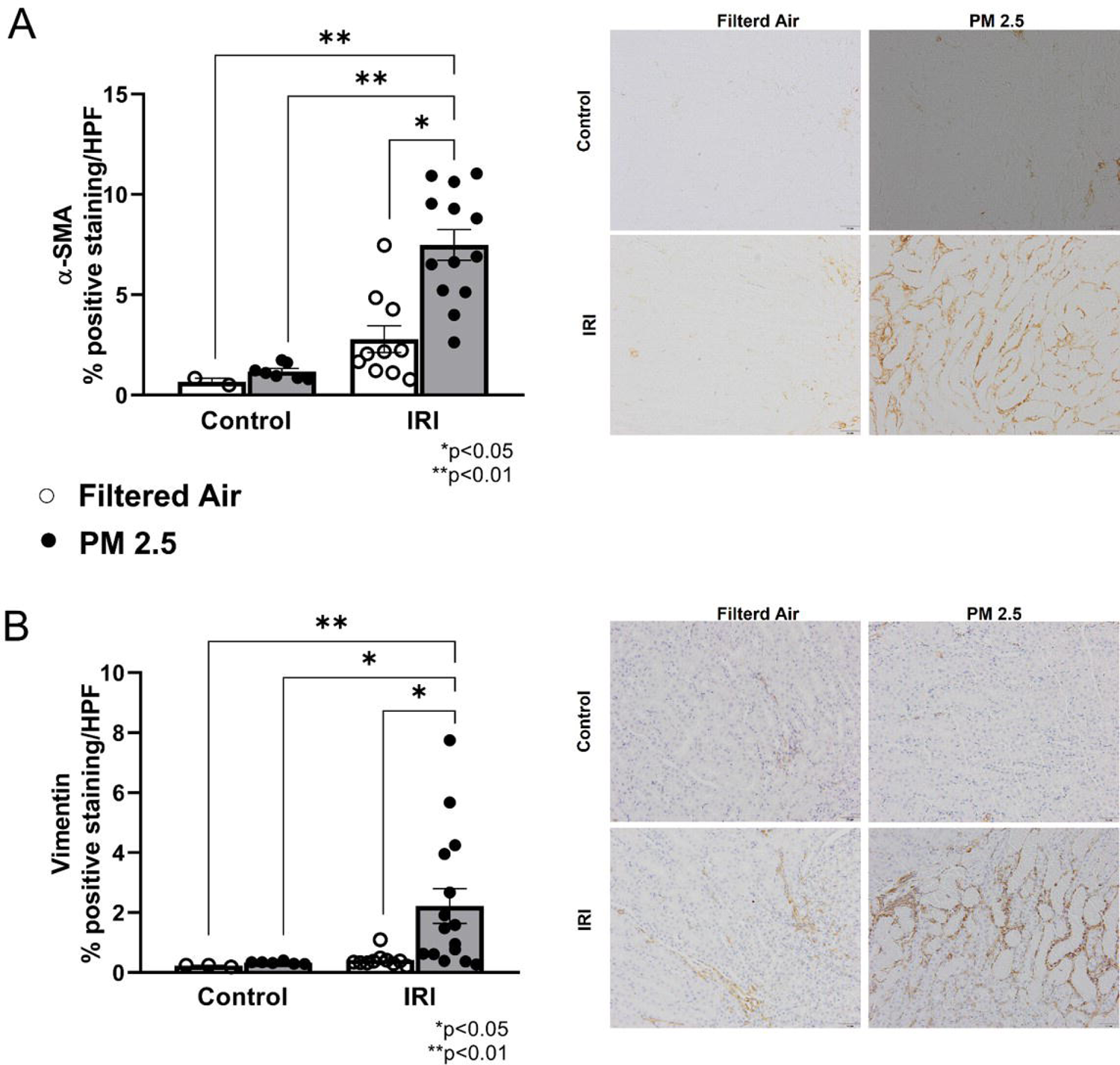
Early hallmarks of renal fibrosis in mice breathing PM2.5 after IRI-induced AKI. (A) Quantification and representative images of kidney tissue sections stained with α-SMA (Kruskal-Wallis test multi comparisons between all groups). (B) Quantification and representative images of kidney tissue sections stained with Vimentin (Kruskal-Wallis test multi comparisons between all groups). White dot/White bar– Filtered Air; Black dot/Gray bar – PM2.5

## DISCUSSION

In this work, through a unique environmentally-controlled setting of PM2.5 exposure, and an experimental model of IRI-induced AKI, we were able to show that PM2.5 aggravates IRI-induced AKI, via enhanced tubular damage, cGAS-STING-mediated inflammation, activation of aging mechanisms and early hallmarks of fibrosis.

Mice were exposed for 12 weeks to a daily target dose of 600 micrograms/m^3^ of PM2.5 before undergoing IRI. This dose was determined based on the accumulated dose that an inhabitant of São Paulo city inhales in 24 hours. Despite being a high dose, it is still relevant for places where PM2.5 is poorly regulated and regularly exceeds the World Health Organization (WHO) air quality guidelines, such as São Paulo, and many other cities around the world. According to the WHO, 9 out of 10 people around the world breathes polluted air^44^.

In such a controlled whole-body exposure system, animals receiving a concentrated stream of PM2.5 and subsequently IRI, presented lower glomerular filtration (measured by creatinine clearance) and reduced ability to concentrate urine, possibly due to more tubular damage. Indeed, the fraction excretion of sodium was increased, as well as markers of tubular damage NGAL and KIM-1. Histologically, this was linked to a profound tubular necrosis. We recognize that we applied here a mild model of IRI (without mortality) which did not exhibit significant changes when compared to control mice, 48 hours after ischemia. However, this mild model proved to be valuable in depicting differences in animals breathing PM2.5 and receiving AKI.

Although previous studies have shown that PM2.5 whole-body exposure in rats or in cultured tubular epithelial cells could induce upregulation of KIM-1 without additional injury^21,22,33^, we were unable to show any change in tubular damage in control mice breathing PM2.5. This could be related to the different animal species used or due to changes in elemental composition of PM2.5 The air quality in São Paulo is influenced by various sources of pollution, with vehicular emissions being the primary contributor to the presence of PM2.5. The extensive use of biofuels, such as gasohol and biodiesel, makes this region a significant case study for comprehending the atmospheric chemistry associated with both fossil fuel and biofuel emissions and may be one of the factors contributing to the different results in terms of kidney damage. PM2.5 in São Paulo is predominantly composed of carbonaceous aerosols, including OC and EC, along with sulfate and nitrate ions^3^. These results align with the analysis we performed of ambient air during the exposure of animals. We have indeed detected high concentrations of PM2.5, OC, and EC, followed by metals. Finally, Nemmar and colleagues^45^ have shown a relationship between PM2.5 exposure and kidney injury, however the particles used in their study appeared to be nanosized. This would suggest that ultrafine particles due to their small size may exert even higher renal toxicity than larger particles. Particle size difference could also explain why our study did not show any difference in tubular damage in control animals exposed to PM2.5.

Proximal tubular epithelial cells (TECs) in the kidney can be considered as sentinels of the immune system^31^. Indeed, damaged TECs during AKI are able to orchestrate an inflammatory response in the kidney. When inflammation exceeds, long-term sequelae after AKI can occur^24^. In control mice, PM2.5 was unable to increase inflammation. Nevertheless, many studies have shown that certain individuals, such as children, elderly and those with pre-exiting diseases, are more susceptible to the effects of PM2.5. In fact, in ischemic kidneys previously exposed to PM2.5, the excessive tubular injury was also associated with an enhanced inflammatory response driven by cGAS-STING. PM2.5 is a significant trigger of systemic inflammation and mitochondria-dysfunction induced oxidative stress. An emerging concept considers mitochondria as a Reactive Oxygen Species (ROS)-signaling integrator and regulators of the innate immune response ^46^.

It is conceivable that after inhalation, deposited particles from the lungs may translocate directly into the circulation, reaching distant organs, including the kidneys. As a proof of concept, nanosized particles seems to have high renal clearance^47^, suggesting that kidney cells come in close contact with these toxins and may therefore contribute to local inflammation and oxidative stress. The kidney plays a vital role in carrying out crucial functions such as eliminating toxic metabolites and drugs from the body. Consequently, it is susceptible to the effects of external toxic substances^48^. This susceptibility stems from the fact that renal tubules, particularly TECs, come in direct contact with drugs during the process of concentration and reabsorption, making them highly vulnerable to environmental toxins. The proximal tubule can be also dramatically affected by mitochondrial damage and inflammation induced during AKI^49^, both known to induce cellular senescence^50^. These changes impact regeneration of the kidney, resulting in accelerated aging and fibrosis^25,51^. p21^Waf^^1/Cip1^, is a protein that binds to CDK2, inhibiting the CDK2–cyclin E complex, promoting cell cycle arrest and senescence^52,53^. Animals breathing PM2.5 with AKI, displayed increased senescence as shown by the enhanced expression of p21 protein in the kidney, but also pro-fibrotic SASP components^54^. p21 is also a universal marker of genotoxic stress and DNA damage induced by air pollutants^55^. We have not found an increase in p21 expression in the control group exposed to PM2.5, however, oxidative stress and DNA damage induced by IRI might be triggering p21 overexpression. Interestingly, a recent work described that mitochondrial dysfunction is able to activate cGAS/STING pathway in vascular smooth muscle cells (VSMCs), leading to premature senescence, resulting in CKD-associated plaque vulnerability^56^. A similar mechanism could take place in TECs during IRI. PM2.5 may induce TECs mitochondrial damage, and activate the cGAS-STING downstream pathway, leading to premature senescence and AKI progression. According with this hypothesis, a recent study found that PM2.5 can accelerate senescence in lung cells via activation of cGAS^57^. Further evidence to support the link between this pro-inflammatory pathway and aging as well as fibrosis is the increased expression of *Cxcl10* gene. *Cxcl10*, which is activated by cGAS-STING, is known to be involved in aging^58^ and its elevated systemic levels are associated with PM2.5 exposure in young adults^59^ and in kidney fibrogenesis^60^.

Indeed we also found that PM2.5 exacerbates inflammation and accelerates aging mechanisms including hallmarks of renal fibrosis. The increased interstitial myofibroblast infiltration observed in the PM2.5+IRI group is considered an early sign of fibrosis, since it precedes the development of renal fibrosis. However, since the sacrifice occurred relatively early (48h after reperfusion) matrix remodeling was not yet evident. These compelling evidences suggest that fibrosis would develop in the animals at a later stage.

PM2.5 has been shown to be involved in different events involved in fibrogenesis in lung^61^ and liver^62^. Exposure to a concentrated stream of ambient PM2.5 led to the development of hepatic fibrosis in mice, regardless of whether they were on a normal chow or high-fat diet. Finally, our group’s collaborators have demonstrated that lupus-prone animals exposed in the same system used in this manuscript, presented an increased cardiac fibrosis, together with heightened levels of IL1β and C3 in the cardiac tissue, indicating an escalation of inflammation^63^.

The reduced expression of the anti-aging protein klotho provided additional evidence for the synergistic impact of PM2.5 and IRI in accelerating kidney aging and senescence^43^. Klotho deficiency is a biomarker of kidney damage that has been associated with aging and senescence in the kidney^43,64^. Environmental pollutants can decrease serum levels of klotho^65^ but the mechanism by which PM2.5 decreases klotho in the kidney remains to be elucidated.

To the best of our knowledge this is the first experimental evidence demonstrating that PM2.5 exposition could aggravate IRI-induced AKI, and may play a role in progression after AKI by accelerating molecular aging mechanisms. The most important question still remains about which component of PM2.5 is toxic for the kidneys and further research is needed to answer this question.

Reducing exposure to PM2.5 can prevent the anticipated burden of kidney disease in the coming years, offering a promising avenue for safeguarding kidney health. This should be a collective effort of governments and individuals.

## ACKNOWLEDGMENTS

This work is supported by a joint consortium grant on healthy aging (www.pmkidney.com) by the Dutch Research Council (NWO) and Fundação de Amparo à Pesquisa do Estado de São Paulo (FAPESP, São Paulo Research Foundation; Grants nos. 2019/19433-0 and 2016/18438-0); ACP, MG and LYUI are the recipients of grants from the Fundação de Amparo à Pesquisa do Estado de São Paulo (FAPESP, São Paulo Research Foundation; Grants nos. 2022/11975-1, 2019/26385-2 and 2018/24171-2 respectively). PS is financially supported by the CSC scholarship from the Chinese government. LA and MFA are the recipients of a grant from the Brazilian Conselho Nacional de Desenvolvimento Científico e Tecnológico (National Council for Scientific and Technological Development; Grants 309683/2021-1 and 306849/2007-0 respectively). AT is financially supported by the NWO-FAPESP joint grant on healthy ageing, executed by ZonMw (no. 457002002).

---

**Table 1.**
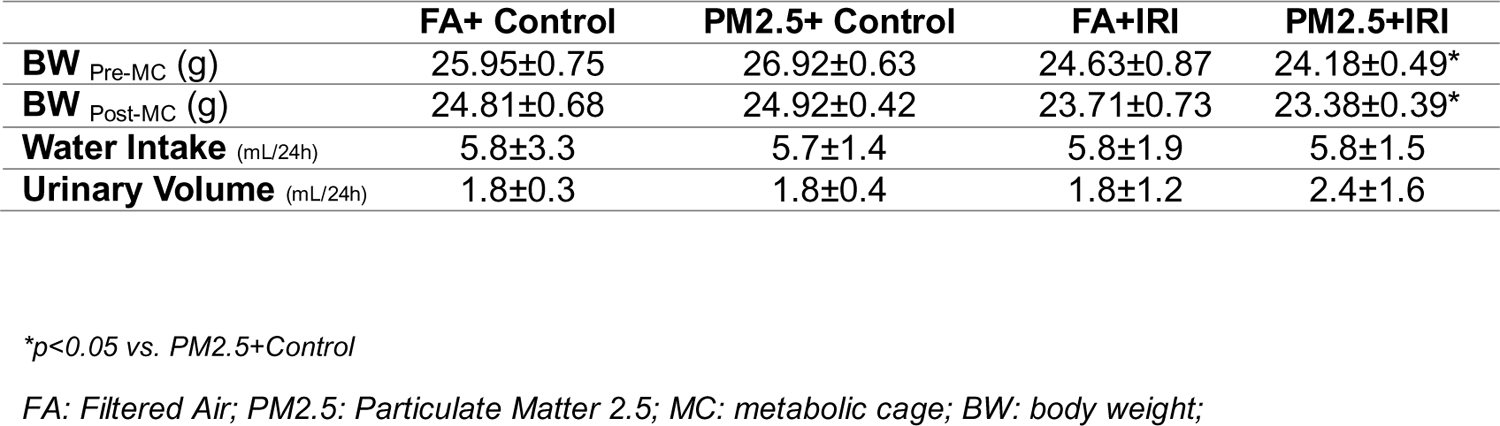
Characteristics of the animals after ischemia/reperfusion surgery.

## REFERENCES

1. Fuller R, Landrigan PJ, Balakrishnan K, et al. Pollution and health: a progress update. The Lancet Planetary Health. 2022;6(6):e535–e547. doi:10.1016/S2542-5196(22)00090-0

2. Thangavel P, Park D, Lee YC. Recent Insights into Particulate Matter (PM2.5)-Mediated Toxicity in Humans: An Overview. International Journal of Environmental Research and Public Health. 2022;19(12). doi:10.3390/ijerph19127511

3. Andrade M de F, Kumar P, de Freitas ED, et al. Air quality in the megacity of São Paulo: Evolution over the last 30 years and future perspectives. Atmospheric Environment. 2017;159:66–82. doi:10.1016/j.atmosenv.2017.03.051

4. De Brito JM, Macchione M, Yoshizaki K, et al. Acute cardiopulmonary effects induced by the inhalation of concentrated ambient particles during seasonal variation in the city of São Paulo. Journal of Applied Physiology. 2014;117(5):492–499. doi:10.1152/japplphysiol.00156.2014

5. Arbex MA, De Souza Conceição GM, Cendon SP, et al. Urban air pollution and chronic obstructive pulmonary disease-related emergency department visits. Journal of Epidemiology and Community Health. 2009;63(10):777–783. doi:10.1136/jech.2008.078360

6. Takano APC, Justo LT, dos Santos NV, et al. Pleural anthracosis as an indicator of lifetime exposure to urban air pollution: An autopsy-based study in Sao Paulo. Environmental Research. 2019;173(November 2018):23–32. doi:10.1016/j.envres.2019.03.006

7. Brook RD, Rajagopalan S, Pope CA, et al. Particulate matter air pollution and cardiovascular disease: An update to the scientific statement from the american heart association. Circulation. 2010;121(21):2331–2378. doi:10.1161/CIR.0b013e3181dbece1

8. Ward-Caviness CK, Weaver AM, Buranosky M, et al. Associations between long-term fine particulate matter exposure and mortality in heart failure patients. Journal of the American Heart Association. 2020;9(6). doi:10.1161/JAHA.119.012517

9. Chen H, Kwong JC, Copes R, et al. Exposure to ambient air pollution and the incidence of dementia: A population-based cohort study. Environment International. 2017;108(May):271–277. doi:10.1016/j.envint.2017.08.020

10. Gatto NM, Henderson VW, Hodis HN, et al. Components of air pollution and cognitive function in middle-aged and older adults in Los Angeles. NeuroToxicology. 2014;40:1–7. doi:10.1016/j.neuro.2013.09.004

11. Hill W, Lim EL, Weeden CE, et al. Lung adenocarcinoma promotion by air pollutants. Nature. 2023;616(7955):159–167. doi:10.1038/s41586-023-05874-3

12. Liu C, Chen R, Sera F, et al. Ambient Particulate Air Pollution and Daily Mortality in 652 Cities. New England Journal of Medicine. 2019;381(8):705–715. doi:10.1056/nejmoa1817364

13. Lelieveld J, Evans JS, Fnais M, Giannadaki D, Pozzer A. The contribution of outdoor air pollution sources to premature mortality on a global scale. Nature. 2015;525(7569):367–371. doi:10.1038/nature15371

14. Bowe B, Xie Y, Li T, Yan Y, Xian H, Al-Aly Z. Particulate matter air pollution and the risk of incident CKD and progression to ESRD. Journal of the American Society of Nephrology. 2018;29(1):218–230. doi:10.1681/ASN.2017030253

15. Bowe B, Xie Y, Li T, Yan Y, Xian H, Al-Aly Z. Estimates of the 2016 global burden of kidney disease attributable to ambient fine particulate matter air pollution. BMJ Open. 2019;9(5). doi:10.1136/bmjopen-2018-022450

16. Chan TC, Zhang Z, Lin BC, et al. Long-term exposure to ambient fine particulate matter and chronic kidney disease: A cohort study. Environmental Health Perspectives. 2018;126(10):1–7. doi:10.1289/EHP3304

17. Bragg-Gresham J, Morgenstern H, McClellan W, et al. County-level air quality and the prevalence of diagnosed chronic kidney disease in the US Medicare population. PLoS ONE. 2018;13(7):1–13. doi:10.1371/journal.pone.0200612

18. Blum MF, Surapaneni A, Stewart JD, et al. Particulate matter and albuminuria, glomerular filtration rate, and incident ckd. Clinical Journal of the American Society of Nephrology. 2020;15(3):311–319. doi:10.2215/CJN.08350719

19. Feng Y, Jones MR, Ahn JYB, Garonzik-Wang JM, Segev DL, McAdams-DeMarco M. Ambient air pollution and posttransplant outcomes among kidney transplant recipients. American Journal of Transplantation. 2021;21(10):3333–3345. doi:10.1111/ajt.16605

20. da Motta Singer J, Saldiva de André CD, Afonso de André P, et al. Assessing socioeconomic bias of exposure to urban air pollution: an autopsy-based study in São Paulo, Brazil. Lancet Regional Health - Americas. 2023;22:100500. doi:10.1016/j.lana.2023.100500

21. Tavera Busso I, Mateos AC, Juncos LI, Canals N, Carreras HA. Kidney damage induced by sub-chronic fine particulate matter exposure. Environment International. 2018;121(August):635–642. doi:10.1016/j.envint.2018.10.007

22. Aztatzi-Aguilar OG, Uribe-Ramírez M, Narváez-Morales J, De Vizcaya-Ruiz A, Barbier O. Early kidney damage induced by subchronic exposure to PM2.5 in rats. Particle and Fibre Toxicology. 2016;13(1):1–20. doi:10.1186/s12989-016-0179-8

23. Xu X, Nie S, Ding H, Hou FF. Environmental pollution and kidney diseases. Nature Reviews Nephrology. 2018;14(5):313–324. doi:10.1038/nrneph.2018.11

24. Tammaro A, Kers J, Scantlebery A, Florquin S. Metabolic flexibility and innate immunity in renal ischemia reperfusion injury: the fine balance between adaptive repair and tissue degeneration. Frontiers in Immunology. 2020;11(July):1346. doi:10.3389/FIMMU.2020.01346

25. Andrade L, Rodrigues CE, Gomes SA, Noronha IL. Acute Kidney Injury as a Condition of Renal Senescence. Cell Transplantation. 2018;27(5):739–753. doi:10.1177/0963689717743512

26. Lawrence J, Wolfson JM, Ferguson S, Koutrakis P, Godleski J. Performance Stability of the Harvard Ambient Particle Concentrator. Aerosol Science and Technology. 2004;38(3):219–227. doi:10.1080/02786820490261735

27. De Oliveira AAF, De Oliveira TF, Dias MF, et al. Genotoxic and epigenotoxic effects in mice exposed to concentrated ambient fine particulate matter (PM2.5) from São Paulo city, Brazil. Particle and Fibre Toxicology. 2018;15(1):1–19. doi:10.1186/s12989-018-0276-y

28. de Barros Mendes Lopes T, Groth EE, Veras M, et al. Pre- and postnatal exposure of mice to concentrated urban PM2.5 decreases the number of alveoli and leads to altered lung function at an early stage of life. Environmental Pollution. 2018;241:511–520. doi:10.1016/j.envpol.2018.05.055

29. Di Domenico M, de Benevenuto SG M, Tomasini PP, et al. Concentrated ambient fine particulate matter (PM2.5) exposure induce brain damage in pre and postnatal exposed mice. NeuroToxicology. 2020;79(May):127–141. doi:10.1016/j.neuro.2020.05.004

30. Leemans JC, Stokman G, Claessen N, et al. Renal-associated TLR2 mediates ischemia/reperfusion injury in the kidney. The Journal of clinical investigation. 2005;115(10):2894–2903. doi:10.1172/JCI22832

31. van der Rijt S, Leemans JC, Florquin S, Houtkooper RH, Tammaro A. Immunometabolic rewiring of tubular epithelial cells in kidney disease. Nature Reviews Nephrology. 2022;18(9):588–603. doi:10.1038/s41581-022-00592-x

32. Maekawa H, Inoue T, Ouchi H, et al. Mitochondrial Damage Causes Inflammation via cGAS-STING Signaling in Acute Kidney Injury. Cell Reports. 2019;29(5):1261–1273.e6. doi:10.1016/j.celrep.2019.09.050

33. Huang X, Shi X, Zhou J, et al. The activation of antioxidant and apoptosis pathways involved in damage of human proximal tubule epithelial cells by PM2.5 exposure. Environmental Sciences Europe. 2020;32(1). doi:10.1186/s12302-019-0284-z

34. Sotty J, Kluza J, De Sousa C, et al. Mitochondrial alterations triggered by repeated exposure to fine (PM2.5-0.18) and quasi-ultrafine (PM0.18) fractions of ambient particulate matter. Environment International. 2020;142(May):105830. doi:10.1016/j.envint.2020.105830

35. Gao M, Liang C, Hong W, et al. Biomass-related PM2.5 induces mitochondrial fragmentation and dysfunction in human airway epithelial cells. Environmental Pollution. 2022;292(PB):118464. doi:10.1016/j.envpol.2021.118464

36. Martens DS, Nawrot TS. Air Pollution Stress and the Aging Phenotype: The Telomere Connection. Current environmental health reports. 2016;3(3):258–269. doi:10.1007/s40572-016-0098-8

37. Ziegler D V., Wiley CD, Velarde MC. Mitochondrial effectors of cellular senescence: Beyond the free radical theory of aging. Aging Cell. 2015;14(1):1–7. doi:10.1111/acel.12287

38. Pardo M, Xu F, Qiu X, Zhu T, Rudich Y. Seasonal variations in fine particle composition from Beijing prompt oxidative stress response in mouse lung and liver. Science of the Total Environment. 2018;626:147–155. doi:10.1016/j.scitotenv.2018.01.017

39. Abbas I, Garçon G, Saint-Georges F, et al. Polycyclic aromatic hydrocarbons within airborne particulate matter (PM2.5) produced DNA bulky stable adducts in a human lung cell coculture model. Journal of Applied Toxicology. 2013;33(2):109–119. doi:10.1002/jat.1722

40. Lee KY, Ho SC, Sun WL, et al. Lnc-IL7R alleviates PM2.5-mediated cellular senescence and apoptosis through EZH2 recruitment in chronic obstructive pulmonary disease. Cell Biology and Toxicology. 2022;38(6):1097–1120. doi:10.1007/s10565-022-09709-1

41. Valentijn FA, Falke LL, Nguyen TQ, Goldschmeding R. Cellular senescence in the aging and diseased kidney. Journal of cell communication and signaling. 2018;12(1):69–82. doi:10.1007/s12079-017-0434-2

42. Tammaro A, Scantlebery AML, Rampanelli E, et al. TREM1/3 Deficiency Impairs Tissue Repair After Acute Kidney Injury and Mitochondrial Metabolic Flexibility in Tubular Epithelial Cells. Frontiers in immunology. 2019;10:1469. doi:10.3389/fimmu.2019.01469

43. Marquez-Exposito L, Tejedor-Santamaria L, Valentijn FA, et al. Oxidative Stress and Cellular Senescence Are Involved in the Aging Kidney. Antioxidants. 2022;11(2):1–19. doi:10.3390/antiox11020301

44. No Title. https://www.who.int/news/item/02-05-2018-9-out-of-10-people-worldwide-breathe-polluted-air-but-more-countries-are-taking-action.

45. Nemmar A, Karaca T, Beegam S, et al. Prolonged pulmonary exposure to diesel exhaust particles exacerbates renal oxidative stress, inflammation and DNA damage in mice with adenine-induced chronic renal failure. Cellular Physiology and Biochemistry. 2016;38(5):1703–1713. doi:10.1159/000443109

46. Banoth B, Cassel SL. Mitochondria in innate immune signaling. Translational Research. 2018;202(3):52–68. doi:10.1016/j.trsl.2018.07.014

47. Miller MR, Raftis JB, Langrish JP, et al. Inhaled Nanoparticles Accumulate at Sites of Vascular Disease. ACS Nano. 2017;11(5):4542–4552. doi:10.1021/acsnano.6b08551

48. Ferguson MA, Vaidya VS, Bonventre J V. Biomarkers of nephrotoxic acute kidney injury. Toxicology. 2008;245(3):182–193. doi:10.1016/j.tox.2007.12.024

49. Stokman G, Kors L, Bakker PJ, et al. NLRX1 dampens oxidative stress and apoptosis in tissue injury via control of mitochondrial activity. The Journal of Experimental Medicine. 2017;214(8):2405–2420. doi:10.1084/jem.20161031

50. Miwa S, Kashyap S, Chini E, von Zglinicki T. Mitochondrial dysfunction in cell senescence and aging. Journal of Clinical Investigation. 2022;132(13):1–9. doi:10.1172/JCI158447

51. Sturmlechner I, Durik M, Sieben CJ, Baker DJ, Van Deursen JM. Cellular senescence in renal ageing and disease. Nature Reviews Nephrology. 2017;13(2):77–89. doi:10.1038/nrneph.2016.183

52. Price PM, Safirstein RL, Megyesi J. The cell cycle and acute kidney injury. Kidney International. 2009;76(6):604–613. doi:10.1038/ki.2009.224

53. Price PM, Safirstein RL, Megyesi J. Protection of renal cells from cisplatin toxicity by cell cycle inhibitors. American Journal of Physiology - Renal Physiology. 2004;286(2 55-2):378–384. doi:10.1152/ajprenal.00192.2003

54. Valentijn FA, Falke LL, Nguyen TQ, Goldschmeding R. Cellular senescence in the aging and diseased kidney. Journal of Cell Communication and Signaling. 2018;12(1):69–82. doi:10.1007/s12079-017-0434-2

55. Rossner P, Binkova B, Milcova A, et al. Air pollution by carcinogenic PAHs and plasma levels of p53 and p21WAF1 proteins. Mutation Research - Fundamental and Molecular Mechanisms of Mutagenesis. 2007;620(1-2):34–40. doi:10.1016/j.mrfmmm.2007.02.020

56. Advanced Science - 2021 - Bi - Mitochondrial Damage-Induced Innate Immune Activation in Vascular Smooth Muscle Cells.pdf.

57. Wu T, Xu S, Chen B, et al. Ambient PM2.5 exposure causes cellular senescence via DNA damage, micronuclei formation, and cGAS activation. Nanotoxicology. 2022;16(6-8):757–775. doi:10.1080/17435390.2022.2147460

58. Yang H, Wang H, Ren U, Chen Q, Chena ZJ. CGAS is essential for cellular senescence. Proceedings of the National Academy of Sciences of the United States of America. 2017;114(23):E4612–E4620. doi:10.1073/pnas.1705499114

59. Pope CA, Bhatnagar A, McCracken JP, Abplanalp W, Conklin DJ, O’Toole T. Exposure to Fine Particulate Air Pollution Is Associated with Endothelial Injury and Systemic Inflammation. Circulation Research. 2016;119(11):1204–1214. doi:10.1161/CIRCRESAHA.116.309279

60. Gao J, Wu L, Zhao Y, Hong Q, Feng Z, Chen X. Cxcl10 deficiency attenuates renal interstitial fibrosis through regulating epithelial-to-mesenchymal transition. Experimental Cell Research. 2022;410(2):112965. doi:10.1016/j.yexcr.2021.112965

61. Guo L, Bai S, Ding S, Zhao L, Xu S, Wang X. PM2.5 Exposure Induces Lung Injury and Fibrosis by Regulating Ferroptosis via TGF-β Signaling. Disease Markers. 2022;2022. doi:10.1155/2022/7098463

62. Zheng Z, Zhang X, Wang J, et al. HHS Public Access. 2016;63(6):1397-1404. doi:10.1016/j.jhep.2015.07.020.Exposure

63. Waked D, Rodrigues ACB, Silva TM, Yariwake VY, Farhat SCL, Veras MM. Effect of chronic exposure to fine particulate matter on cardiac tissue of NZBWF1 mice. International Journal of Experimental Pathology. 2023;(February):177–187. doi:10.1111/iep.12473

64. Hu MC, Shi M, Zhang J, Quĩones H, Kuro-O M, Moe OW. Klotho deficiency is an early biomarker of renal ischemia-reperfusion injury and its replacement is protective. Kidney International. 2010;78(12):1240–1251. doi:10.1038/ki.2010.328

65. Yao Y, He G yan, Wu X juan, et al. Association between environmental exposure to perchlorate, nitrate, and thiocyanate and serum α-Klotho levels among adults from the National Health and nutrition examination survey (2007–2014). BMC Geriatrics. 2022;22(1):1–8. doi:10.1186/s12877-022-03444-2

