## Supplemental Table 1 for "Air Pollution Aggravates Renal Ischemia-Reperfusion-Induced Acute Kidney Injury"

**Supplemental Table 1: Concentration of PM2.5 and composition**

| **Pollutant** | **ug/m^3^** |
| --- | --- |
| PM2.5 | 29.8±11 |
| Organic Carbon (OC) | 9.67 ±4 |
| Elemental Carbon (EC) | 3.79±2.1 |
| Magnesium (Mg) | 0.18±0.1 |
| Aluminum (Al) | 0.27±0.2 |
| Potassium (K) | 0.27±0.2 |
| Titanium (Ti) | 0.01±0.01 |
| Vanadium (V) | 0.001±0.001 |
| Chromium (Cr) | 0.002±0.002 |
| Manganese (Mn) | 0.005±0.003 |
| Iron (Fe) | 0.19±0.1 |
| Nickel (Ni) | 0.0007±0.0005 |
| Copper (Cu) | 0.05±0.02 |
| Zinc (Zn) | 0.03±0.03 |
| Arsenic (As) | 0.0003±0.0002 |
| Selenium (Se) | 0.0008±0.0007 |
| Rubidium (Rb) | 0.001±0.0009 |
| Strontium (Sr) | 0.002±0.001 |
| Molybdenum (Mo) | 0.0015±0.001 |
| Silver (Ag) | 6.25E-05±3.46E-05 |
| Tin (Sn) | 0.005±0.005 |
| Barium (Ba) | 0.011±0.008 |
| Lead (Pb) | 0.008±0.008 |
| Bismuth (Bi) | 0.0001±8.6E-05 |
| Uranium (U) | 4.01E-05±2.82E-05 |
| Sodium (Na)* | 2.54±0.3 |
| Ammonium (NH_4_)* | 0.41±0.4 |
| Potassium (K) * | 0.21±0.1 |
| Calcium (Ca_2_)* | 1.32±0.5 |
| Chlorine (Cl)* | 0.29±0.2 |
| Nitrate (NO_3_)* | 1.9±1.2 |
| Sulfate (SO_4_)* | 2.09±1.5 |

Membrane filters sampled at the University of Sao Paulo (campus Butantã) were analyzed for the identification of PM2.5 mass and composition of trace elements through inductively coupled plasma mass spectrometry (ICPMS), ion chromatography and mass gravimetric analysis.

* Measured with ion chromatography; data are expressed as mean ± SD
